# A Back-Door Insights into the modulation of Src kinase activity by the polyamine spermidine

**DOI:** 10.1101/2023.01.15.524120

**Authors:** Sofia Rossini, Marco Gargaro, Giulia Scalisi, Elisa Bianconi, Sara Ambrosino, Eleonora Panfili, Claudia Volpi, Ciriana Orabona, Antonio Macchiarulo, Francesca Fallarino, Giada Mondanelli

## Abstract

Src is a protein tyrosine kinase commonly activated downstream of transmembrane receptors and plays key roles in cell growth, migration and survival signaling pathways. In conventional dendritic cells (cDCs), Src is involved in the activation of the non-enzymatic functions of indoleamine 2,3-dioxygenase 1 (IDO1), an immunoregulatory molecule endowed with both catalytic activity and signal transducing properties. Prompted by the discovery that the metabolite spermidine confers a tolerogenic phenotype on cDCs that is dependent on both the expression of IDO1 and the activity of Src kinase, we here investigated the spermidine mode of action. We found that spermidine directly binds Src in a previously unknown allosteric site located on the backside of the SH2 domain and thus acts as a positive allosteric modulator of the enzyme. Besides confirming that Src phosphorylates IDO1, here we showed that spermidine promotes the protein-protein interaction of Src with IDO1. Overall, this study may pave the way toward the design of allosteric modulators able to switch on/off the Src-mediated pathways, including those involving the immunoregulatory protein IDO1.

## Introduction

Besides being intermediates in metabolic reactions, metabolites can serve as intracellular and intercellular signals (1). Indeed, by interacting with specific molecular partners, soluble mediators can trigger a series of molecular events critical for cell fitness and adaptation. Metabolites binding to either the active site or the allosteric pocket – i.e., that different from the catalytic site – of enzymes are among the best-characterized interactions that modulate protein activity as well as the assembly and function of multiprotein complexes (2–4).

The naturally occurring polyamines (i.e., putrescine, spermidine and spermine) are organic cations derived from the decarboxylation of L-ornithine, which is generated by the arginase 1 from L-arginine (5,6). The conversion of L-ornithine into putrescine is catalyzed by the rate-limiting enzyme ornithine decarboxylase 1, followed by two specific synthases that sequentially give rise to spermidine and spermine (7). These metabolites are protonated at physiological pH levels, allowing them to interact with negatively charged macromolecules, including nucleic acids, proteins, and phospholipids. Given their structure, polyamines indeed modulate several cellular processes, ranging from cell growth and proliferation to immune system function (8,9). As a matter of the fact, alteration of polyamines intracellular content is associated with the occurrence of several tumors, including prostate, breast, and colon cancers, for which polyamines are considered as biomarkers (10–12). Among polyamines, spermidine has recently gained much more attention as player of immune regulation and in age-related disorders, such as cardiac hypertrophy and memory impairment (13–18). Spermidine exerts a protective role in mouse experimental models of autoimmune diseases, such as multiple sclerosis and psoriasis, by activating the Forkhead box protein O3 (FOXO3) pathway and thus suppressing the production of inflammatory cytokines tumor necrosis factor (TNF)-α and interleukin (IL)-6 (14). Moreover, spermidine is able to reprogram mouse conventional dendritic cells (cDCs) toward an immunoregulatory phenotype *via* Src kinase-dependent phosphorylation of indoleamine 2,3-dioxygenase 1 (IDO1) (19). However, the exact mechanism of Src activation by spermidine remains to be elucidated.

The non-receptor tyrosine kinase Src is the representative of a family of structure-related kinases initially discovered as a proto-oncogene regulating critical cellular functions (20). Src activation mainly occurs downstream of multiple transmembrane receptors, including epidermal growth factor receptor (EGF-R), fibroblast growth factor receptor (FGF-R), and insulin-like growth factor-1 receptor (IGF-1R). Indeed, a dysregulated Src activity has been associated with tumor growth and metastasis, inflammation-mediated carcinogenesis, and therapeutic resistance to traditional antineoplastic drugs (21–26). The induction of Src kinase activity can also occur following Aryl hydrocarbon Receptor (AhR) activation, whose conformational changes favor the Src-AhR disjunction, allowing the former to phosphorylate its downstream partner IDO1 and thereby promote the generation of an immunoregulatory milieu (27).

In addition to the kinase domain, Src possesses an N-terminal Src homology-4 (SH4) domain, a unique domain, an SH3 domain, an SH2 domain, an SH2-kinase linker domain, and a C-terminal autoregulatory motif (25). The SH2 and SH3 modules serve in protein-protein interactions that are essential for the regulation of kinase activity and signaling function. Specifically, the SH2 domain contains two distinct binding pockets. The first one has a conserved arginine residue that binds a phosphotyrosine (pY) residue presented by the protein substrate, whereas the second pocket binds the residue that is three positions C-terminal of pY (pY+3), contributing to the specificity in ligand protein recognition.

The autoregulatory function of the kinase occurs through intramolecular interactions that stabilize the catalytically inactive conformation of Src, in which the SH2 domain binds to a pY located at position +535. Accordingly, binding of ligand proteins to the SH2 domain displaces intramolecular contacts and promotes the catalytic activation of the Src kinase, which is characterized by the phosphorylation of a tyrosine residue in the activation loop (Y424). Given the crucial role of the non-catalytic domains in modulating Src kinases activity, efforts have been made to develop drug-like modulators of the SH2 and SH3 domains. Small peptidomimetics destabilize the closed conformation and thus promote the kinase activation through the binding of SH3 and/or SH2 domains (28,29). Alternatively, modulators of Src kinases able to reinforce the intramolecular interactions have proven to allosterically inhibit the enzyme activity (30).

Prompted by the finding that spermidine triggers the immunosuppressive IDO1 signaling in cDCs (19), here we investigated the molecular relationship between that polyamine, Src kinase and IDO1. We found that spermidine (*i*) activates Src kinase with an allosteric mechanism; (*ii*) binds directly Src kinase at a previously unknown allosteric site; (*iii*) favors the association of IDO1 and Src kinase.

## Results

### Spermidine causes allosteric activation of the kinase activity of Src

The activation of Src kinase mainly occurs downstream of multiple transmembrane and intracellular receptors (such as AhR) as well as protein tyrosine phosphatases (27,31,32). In cDCs, it has been demonstrated that a small molecule, namely spermidine, activates Src with a still undefined mechanism (19). To figure out whether a direct activation would occur, we assayed spermidine against purified recombinant human Src (rhSrc) protein. After 30 minutes of incubation, a luminescent assay was used to measure the ADP released by the kinase. Results showed that spermidine activated rhSrc with a half-maximal effective concentration (EC50) of 106.4 nM ± 13.4 (**Fig 1A**). To confirm the modulation of the kinase also in living cells, we resorted to immunoblot analysis of phosphorylated Src at the tyrosine Y424 as sign of kinase activation. SYF cells (i.e., fibroblast null for Src family kinases, Src, Yes, and Fyn (33)) were stably transfected with vector encoding for Src kinase and then treated with increasing concentration of spermidine. Results showed that the metabolite promoted Src phosphorylation with an EC50 of 6.4 μM ± 0.6 (**Fig 1B, 1C**).

**Figure 1.**
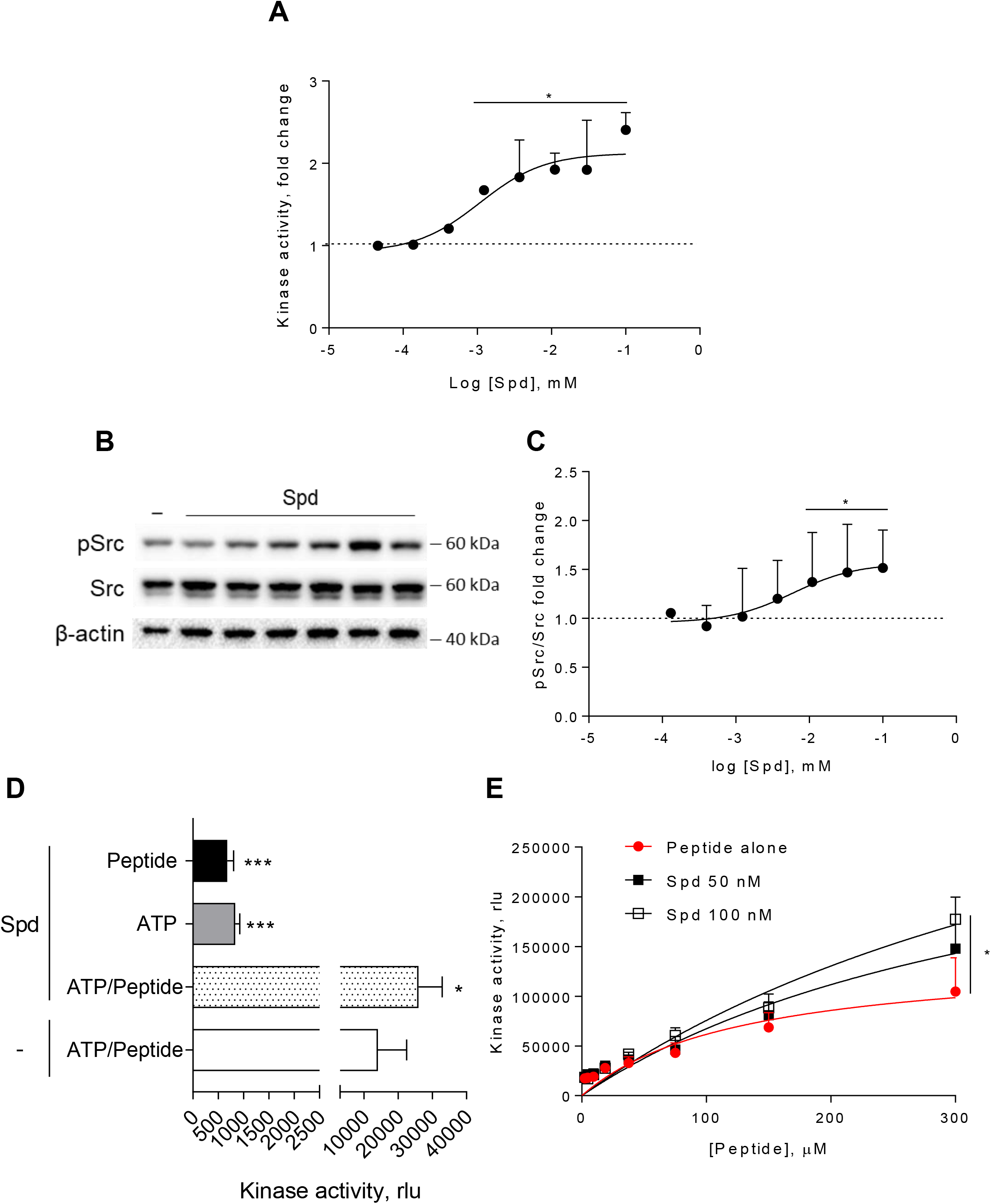
Spermidine enhances the activity of Src kinase in ATP-independent manner. (**A**) Enzymatic activity of rhSrc in the presence of ATP (10 μM), synthetic peptide (100 μM) and increasing concentration of spermidine (45 nM to 100 μM). ADP-Glo™ Kinase Assay (Promega) was used to detect the activity. Results are shown as fold change vs untreated samples (fold change = 1, dotted line). Spermidine EC50=106,4 nM ± 13,4. (**B**) Immunoblot analysis of phosphorylated (pSrc) and total Src protein level evaluated in cell lysates from SYF cells reconstituted with vector coding for wild-type Src and then treated with increasing concentration of spermidine (130 nM to100 μM). Actin expression was used as normalizer. One representative immunoblot of three is shown. (**C**) pSrc/Src ratio of scanning densitometry analysis of three independent immunoblots. Data (mean ± SD) are reported as fold change of samples treated with spermidine relative to untreated cells (fold change = 1, dotted line). Spermidine EC50=6,4 μM ± 0,6. (**D**) Enzymatic activity of rhSrc in the presence of spermidine, with or without ATP and peptide substrate. (**E**) Enzymatic activity of rhSrc in the presence of fixed concentrations of spermidine and increasing concentration of peptide substrate. Data were analyzed with one-way ANOVA followed by post-hoc Bonferroni test. *p < 0.05, ***p < 0.001.

To get insights into the mechanism of action of spermidine, we measured the intrinsic activity of the polyamine in the absence of either ATP or the synthetic peptide. Results showed that spermidine did not activate Src in the absence of either ATP or peptide (**Fig 1D**), while it promoted the production of ADP when the substrate is also present, ruling out any competition for the same site. As this profile was compatible with an allosteric modulation, we incubated rhSrc with different concentration of ATP or peptide. In the presence of fixed amount of spermidine and increasing concentration of the peptide, the maximum rate of Src kinase activity (Vmax) increased, while the affinity (Km) for the substrate was not affected (**Fig 1E**). On the contrary, in the presence of different concentration of ATP, spermidine did not modify neither the efficacy nor the affinity of Src kinase (**Supplementary Fig S1**). Such a kinetic profile is consistent with a non-ATP competition, suggesting that spermidine allosterically activates the kinase activity of Src.

### Spermidine binds to a negatively charged pocket in SH2 domain of Src kinase

In the inactive state, Src assumes a closed conformation with the SH3 domain bound to the SH2-kinase linker and the SH2 domain bound to the tyrosine phosphorylated tail (**Figure 2A**). Using the experimental available structure of SH2 domain (PDB ID: 2JYQ) and electrostatic potential calculations, we characterized key structural and electrostatic elements of the SH2 domain involved in ligand protein recognition (**Fig 2B**). A positive electrostatic potential was observed in the region of the pY binding site (R183 and H209 residues, **Fig 2B**), whereas a stretch of surface endowed with a strong negative electrostatic potential was observed on the backside of the pY binding site as delimited by glutamate residues E155 and E174 (**Fig 2B**), suggesting the existence of a putative allosteric site. Of note, by the alignment of amino acid sequences, we identified that such residues were conserved in Src protein of human, murine, chicken and rat (**Supplementary Fig S2**), further supporting potential functional role for this allosteric site.

**Figure 2.**
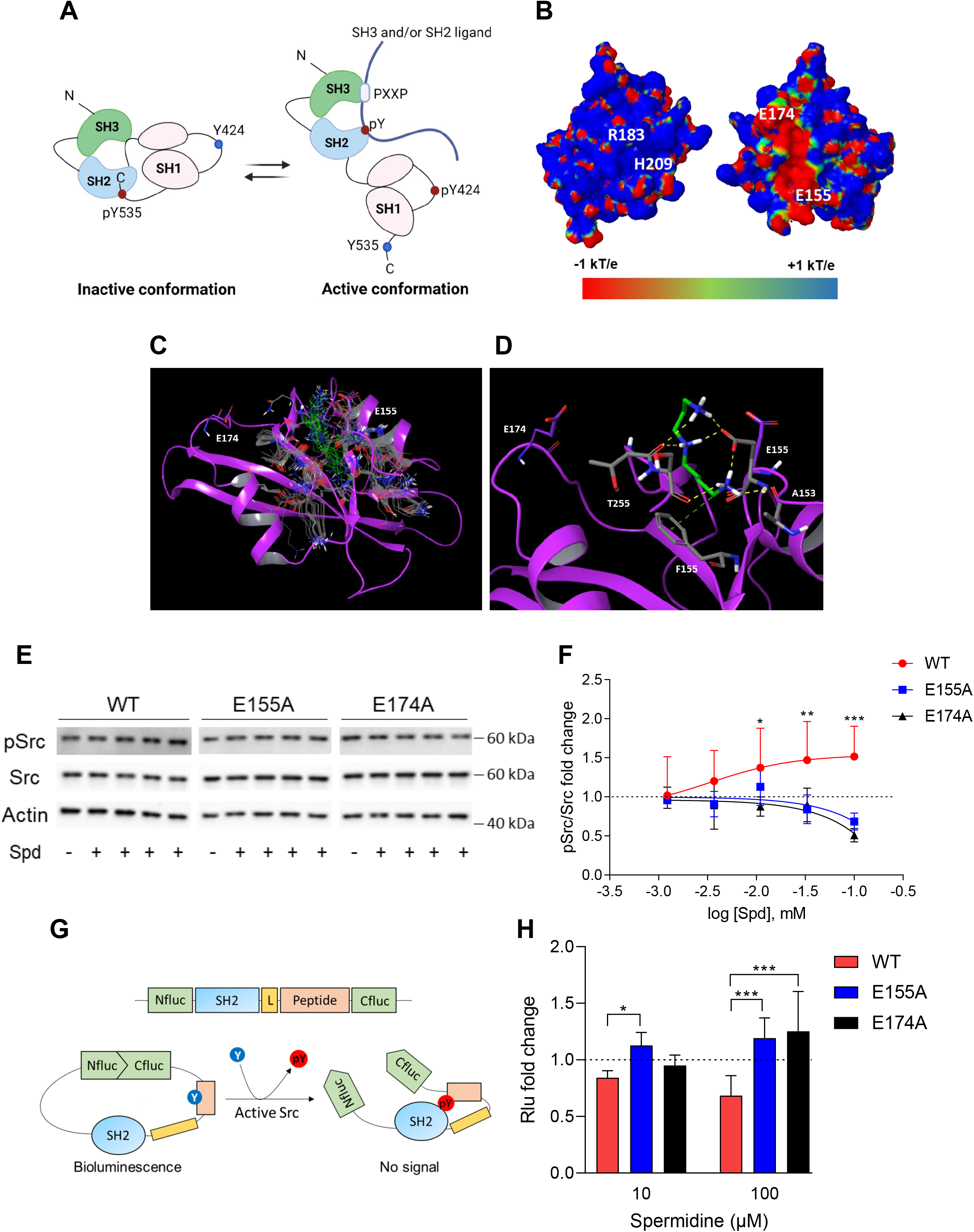
Spermidine binds to an allosteric site located in the SH2 domain of Src kinase. (**A**) Schematic representation of the Src domains and kinase activation. The catalytic activation of the enzyme is characterized by the phosphorylation of the Y424 (pY424) in the activation loop. Created with BioRender.com. (**B**) Electrostatic potential surface of the Src SH2 domain showing the pY binding site (R182, H209) and the putative allosteric site for the endogenous polyamine as delimited by the glutamate residues (E155 and E174). (**C**) Overlay of docking solutions of spermidine into the shallow cavity of Src kinase (poses #1-18, Appendix Table S1). E155 and E174 residues are shown with magenta carbon-atoms. Induced-fit conformations of side chains of residues shaping the cavity are shown with grey carbon-atoms according to each docking solution. Conformations of spermidine according to each docking solution are shown with green carbon-atoms. The Src SH2 domain is shown with magenta cartoon depicting the secondary structure. (**D**) Best energy-scored solution of the binding mode of spermidine into the allosteric pocket of Src (pose #1, Appendix Table S1). E155 and E174 are shown with magenta carbon-atoms. Interacting residues and spermidine are shown with grey and green carbon-atoms, respectively. Hydrogen bond interactions are shown with yellow dashed lines, while the π-cation interaction is reported with green dashed line. (**E**) Immunoblot analysis of phosphorylated (pSrc) and total Src protein level in cell lysates from SYF cells either reconstituted with vector coding for wild-type Src (WT) or Src mutated at glutamate 155 or 174 with alanine (E155A; E174A). Cells were then exposed to increasing concentration of spermidine (130 nM to 100 μM). Actin expression was used as normalizer. One representative immunoblot of three is shown. (**F**) Activation of Src kinase in SYF cells treated as in (**E**) and measured as pSrc/Src ratio of scanning densitometry analysis of three independent immunoblots. Results (mean ± SD) are reported as fold change of samples treated with spermidine relative to untreated cells (fold change = 1, dotted line). (**G**) Schematic representation of the reporter functions. In the presence of active Src kinase, the phosphorylation of Src peptide results in its intra-molecular interaction with the SH2 domain that prevents the complementation of split luciferase fragments and generates a reduced bioluminescence activity. In the absence of Src activation, the N- and C-terminal luciferase domains are reconstituted and thus the bioluminescent activity is restored. (**H**) Measurement of luminescent signal in SYF cells co-expressing the reporter and the wild-type Src or its mutants (E155A and E174A), and then exposed to spermidine (at 10 µM and 100 µM). Results are reported as fold change of bioluminescent signal in stimulated cells as compared to untreated samples. Data (**F, H**) were analyzed with 2-way ANOVA followed by post-hoc Bonferroni test. *p < 0.05, **p < 0.01, ***p < 0.001.

A docking study was carried out to investigate the binding mode of spermidine into the allosteric site of Src SH2 domain. As a result, n.18 solutions were obtained showing a conserved binding mode located in a shallow cavity close to E155 and shaped by A153, F155 and T255 (**Fig 2C**). According to the top scored solution (**Supplementary Table S1**; **Fig 2D**), the first primary amine group interacts by an electrostatic enforced hydrogen bond with E155, the secondary amine group forms electrostatic enforced hydrogen bond with E155 and the carbonyl group of T255, the other primary amine group makes hydrogen bonds with the side chain of E155 and the carbonyl group of A153 while engaging the aromatic ring of F155 through a specific π-cation interaction (34).

To experimentally confirm the proposed spermidine binding site, we resorted to mutagenesis experiments by substituting the glutamate residues 155 or 174 of murine Src into alanine (E155A; E174A). SYF cells were thus stably transfected with vectors coding for the mutated Src (i.e., Src E155A and Src E174A) and wild-type Src (WT) (**Supplementary Fig S3A**). To validate the functional equivalence of Src mutants, cells were exposed to lysophosphatidic acid (LPA), a stimulus known to activate Src kinase downstream the LPA2 receptor in SYF cells (35). Results indicated that Src activity is induced by LPA as measured by the phosphorylation of the Y424, independently of the mutation at the putative allosteric site (**Supplementary Fig S3B**). On evaluating the activation of Src by spermidine, we found that the mutation of the glutamate residues abrogated the kinase activation (**Fig 2E, 2F**). The split-luciferase fragment complementation assay confirmed that E155 and E174 are key anchoring points for spermidine binding. Specifically, SYF cells expressing Src WT or mutant were stably transfected with a bioluminescent reporter that contains the SH2 domain and the Src consensus substrate peptide between the amino-(Nluc) and carboxyl-(Cluc) terminal domains of the Firefly luciferase molecule (**Fig 2G**) (36). When the endogenous Src is active, the tyrosine residue of the consensus peptide is phosphorylated and interact with the docking pocket of the SH2 domain. This creates a steric hindrance that prevents the reconstitution of a functional luciferase, resulting in a reduction of bioluminescent signal (**Fig 2G**). Cells co-expressing Src and the reporter were thus exposed to spermidine and the luminescent signal was measured. Results demonstrated that the bioluminescence decreased when spermidine is applied only in cells ectopically expressing wild-type Src (**Fig 2H**).

Overall, these data suggested the presence of a previously unknown allosteric site on the backside of Src SH2 domain as defined by the glutamate residues at position 155 and 174. Spermidine, by means of ionic and hydrogen bond interactions between its protonated amino groups and residues of the shallow anionic site on the SH2 domain, directly associates with and activates Src kinase. It is worth noting that no direct interaction was observed between spermidine and E174 in the docking study. This may be ascribed to the limit of the scoring function in identifying a binding mode engaging E174 among resulting solutions, or to an indirect role of such residue in promoting long range electrostatic interactions to accomplish the molecular recognition of the cognate ligand into the allosteric site.

### Spermidine promotes the Src-dependent tyrosine phosphorylation of IDO1 and their interaction

Among the proteins phosphorylated by Src, the immunometabolic enzyme IDO1 is worthy of note (19,27). Indeed, aside metabolizing the amino acid tryptophan, IDO1 is endowed with non-enzymatic properties (31,37–39). The latter relies on the presence of two ITIMs (immunoreceptor-tyrosine based inhibitory motif) that can be phosphorylated in response to immunomodulatory stimuli, such as TGF-β, L-kynurenine and spermidine (19,27,38). However, the exact molecular mechanism and the role of spermidine have never been explored. To confirm that Src can phosphorylate IDO1, SYF cells were reconstituted with vectors coding for wild-type Src and IDO1, either alone or in combination, and then were exposed to spermidine. Results from immunoblot demonstrated that the co-precipitated IDO1 is tyrosine phosphorylated by Src and that the polyamine increases the phosphorylation (**Fig 3A**). To further confirm that spermidine could promote the IDO1 phosphorylation by accelerating the reaction velocity, an in vitro kinase assay was performed using purified Src and IDO1 protein. By detecting phosphotyrosine residues with a specific antibody, we found that IDO1 was phosphorylated in a time dependent manner (**Fig 3B, 3C**). Moreover, in the presence of spermidine, Src quicker phosphorylated IDO1, as demonstrated by the 2-fold increase of the relative velocity (**Fig 3D**). To figure out whether the IDO1 phosphorylation was a direct effect through physical interaction with Src, SYF cells reconstituted with wild-type Src and IDO1 were exposed to spermidine for different length of time. Co-immunoprecipitation followed by immunoblot studies demonstrated that when cells were treated with spermidine for 60 minutes, IDO1 was found in a complex with Src (**Fig 3E**). The specific IDO1-Src interaction was confirmed *in situ* by the proximity ligation assay (**Fig 3F, 3G**). Accordingly, spermidine treatment induced the Src-IDO1 interaction in SYF cells reconstituted with wild-type Src, but not with the E155A or E174A mutant form of the kinase (**Fig 3F, 3G**).

**Figure 3.**
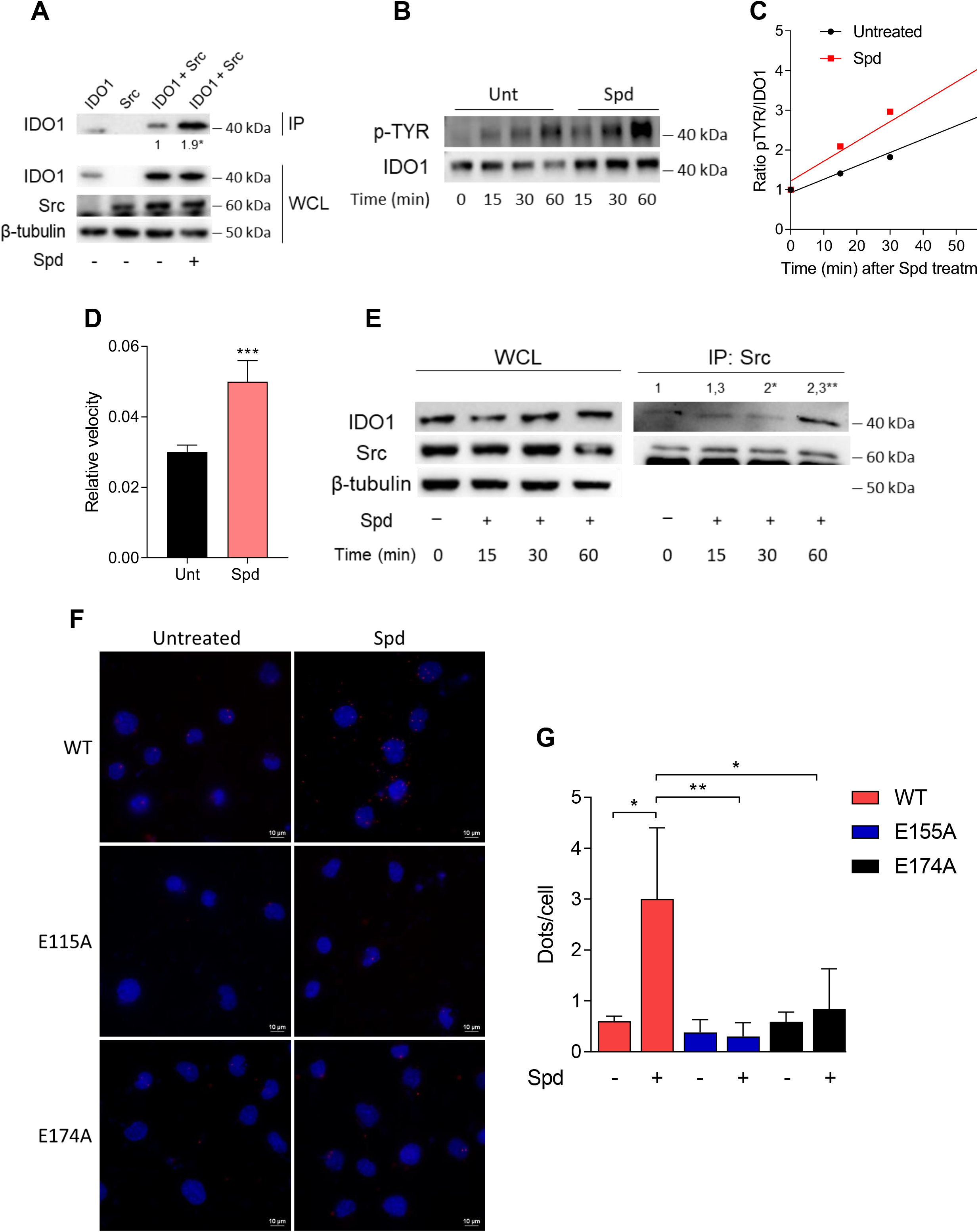
Spermidine triggers the phosphorylation of IDO1 by Src kinase and the complex formation. (**A**) Immunoprecipitation with anti-phosphotyrosine antibody from SYF cells reconstituted with vectors coding for Src and IDO1 and then treated with spermidine (100 μM) for 60 minutes. Cells transfected with vectors coding for either Src or IDO1 were used as control. The detection of IDO1, Src and β-tubulin was performed by sequential immunoblotting with specific antibodies. Whole-cell lysates (WCL) was used as control of protein expression. One representative immunoblot of three is shown. IDO1/pTYR ratio is measured by densitometric quantification of the specific bands and is expressed relative to untreated cells. (**B**) Continuous in vitro kinase assay with rhIDO1 (300 ng) and rhSrc (50 ng) followed by immunoblot analysis with anti-phosphotyrosine and anti-IDO1 specific antibodies. The reaction was carried out for the indicated time, in either the presence or absence of spermidine. One representative immunoblot of three is shown. (**C**) pTYR/IDO1 signals were calculated by densitometric quantification of the specific bands. Data were plotted over incubation time of the kinase reaction and the slopes (relative velocity) of linear fits were calculated. (**D**) The relative velocity of the kinase reaction in either the presence or absence of spermidine from three independent experiments is shown. (**E**) Immunoprecipitation of Src from SYF cells reconstituted with Src and IDO1, and then treated with spermidine for the indicated time. The detection of IDO1, Src and β-tubulin was performed by sequential immunoblotting with specific antibodies. Whole-cell lysates (WCL) was used as control of protein expression. One representative immunoblot of three is shown. IDO1/Src ratio is calculated by densitometric quantification of the specific bands and is reported as fold change against untreated cells. (**F**) The *in situ* proximity ligation assay between IDO1 and Src in SYF cells reconstituted with wild-type Src or the mutant forms and treated as in (**A**). Red spots indicate a single IDO1/Src interaction; scale bars, 10 µm. One representative experiment of three is shown. (**G**) Quantification of the interactions detected by proximity ligation assay using ImageJ. Results are reported as function of the number of cells. Data (mean ± SD) in (**A, E, G**) were analyzed with one-way ANOVA followed by post-hoc Bonferroni test. *p < 0.05, **p < 0.01. Data (mean ± SD) in (**D**) were analyzed with unpaired student t-test. ***p < 0.001.

As a whole, these results suggested that spermidine not only accelerates the Src-mediated phosphorylation of IDO1, but also promotes the formation of Src-IDO1 complex.

## Discussion

The non-receptor tyrosine kinase Src is the representative of a family of structure-related enzymes involved in several signaling pathways regulating key cellular processes as well as immune responses (21,40). Much relevant literature correlates dysregulated Src kinase activity with cancer and thus extensive efforts have been made to develop small molecules kinase inhibitors. Currently approved kinase inhibitors are compounds that reversibly bind the catalytic site and thus compete with the ligand (i.e., ATP) (25). As the ATP-binding cleft is structurally well-conserved among kinases, these inhibitors are poorly selective. Moreover, their chronic usage is frequently associated with acquired drug resistance that ultimately limits patients’ compliance and the therapeutic success. For instance, the FDA-approved Dasatinib and Bosutinib inhibit more than 30 kinases, and thus are not suitable for probing Src-dependent pharmacology (41). Saracatinib is another example of small molecule that interacts with the ATP-binding pocket. Although more selective than Dasatinib, it potently inhibits EGFR as well (42). In addition to competitive Src inhibitors, an emerging pharmacological modality – known as targeted covalent inhibitors (TCIs) – has been pursued at the preclinical level for blocking Src kinase activity (43). However, the promiscuity of molecules interacting with the ATP pocket has moved the interest toward the development of alternative strategies for more effective and less-toxic inhibitors.

The peculiarity of the Src protein, as well as of other tyrosine kinases, is its structural plasticity, i.e., the capability to adopt distinct conformations due to intrinsic dynamic properties (44). The activation state of this protein kinase is indeed dictated by dynamic intramolecular interactions between the SH3, SH2 and kinase domains. The SH2 domain plays a key role in both autoregulating Src kinase activity and in recruiting the protein ligand. Specifically, the tyrosine residue at position +535, when phosphorylated, interacts with the SH2 domain and stabilizes a restrained catalytically inactive conformation of Src (45). Accordingly, binding of ligand proteins to the SH2 domain displaces intra-molecular contacts and promotes the catalytic activation of the Src kinase. This activating event is mostly driven by a dynamic breakage and formation of electrostatic interactions that involve salt bridges and hydrogen bonds. Guided by the dynamic nature of the kinase, allosteric modulation has been proposed as pharmacological approach to target the activity of Src kinase. Allosteric molecules do not possess intrinsic efficacy, but instead modulate – either positively or negatively – the activity of orthosteric agonists. Moreover, being less conserved among kinases, the allosteric hotspots ensure greater drug selectivity. Targetable allosteric pockets have been identified for few kinases as reported for Hck, Lyn, Aurora A kinase, and Bcr-Abl (30,46–48). In addition, modulators of the SH2 and SH3 domains – either peptidomimetics or small molecules – have been developed as chemical tools modifying the conformation and thus both the enzymatic and non-enzymatic functions of Src, the latter including protein-protein interactions and intracellular localization (28,29,47,49,50).

Metabolites are chemicals that do not merely take place in the metabolic reactions, but are also involved in inter- and intra-cellular communications, energy production, macromolecule synthesis, post-translational modifications, and cell survival (51–56). In accordance, the enzymes responsible for their production are considered central regulators of the function of cells, including immune cells (57–61). IDO1 is the prototype of such metabolic enzymes acting at the forefront of immune responses. Thanks to its catalytic activity as well as nonenzymatic function (relying on the phosphorylation of its ITIMs), IDO1 is a tiebreaker of tolerance and immunity (58,62,63). Prompted by the finding that spermidine (i.e., a natural occurring polyamine) can reprogram murine cDCs toward an immunoregulatory phenotype *via* the Src kinase-dependent induction of the IDO1 signaling (19), we here demonstrated that the polyamine behaves as a positive allosteric modulator of Src by increasing the maximum rate of enzyme activity. Indeed, electrostatic potential calculation studies on the SH2 domain identified a surface endowed with a negative electrostatic potential on the back side of the pY binding site, as delimited by glutamate residues E155 and E174. By its protonated amino groups, spermidine interacts with the anionic head of E155 on the SH2 domain, directly associates with and activates Src kinase. As a matter of the fact, the site-directed mutagenesis of the glutamate residues with uncharged amino acids abrogates the spermidine-mediated activation of Src kinase. It is noteworthy of mention that polyamines are not new in the field of allosteric modulation, as they modify the activity of ionotropic N-methyl-D-aspartate receptor (NMDAR, a receptor for glutamate) by both increasing the affinity of NMDAR for the co-agonist glycine and relieving the tonic proton inhibition of the receptor (64), further supporting the spermidine mode-of-action.

Besides confirming that Src phosphorylates IDO1, and that the polyamine accelerates the enzyme kinetic, here we showed that spermidine promotes the interaction of Src with IDO1 protein (**Fig 4**). Our data provided evidence that an endogenous metabolite, when present at specific concentrations, can directly activate Src kinase without requiring a membrane receptor. By acting on the backside of the SH2 domain, that is the domain responsible for the substrate binding, spermidine not only modulates the catalytic activity, but also affects the scaffold function of Src in organizing transducing signaling complexes – as those with IDO1 - which could be relevant in many diseases. Thus, from a therapeutic perspective, our results provide the proof of principle for the development of molecules that can modulate the kinase activity and the nonenzymatic functions of Src and IDO1 at once.

**Figure 4.**
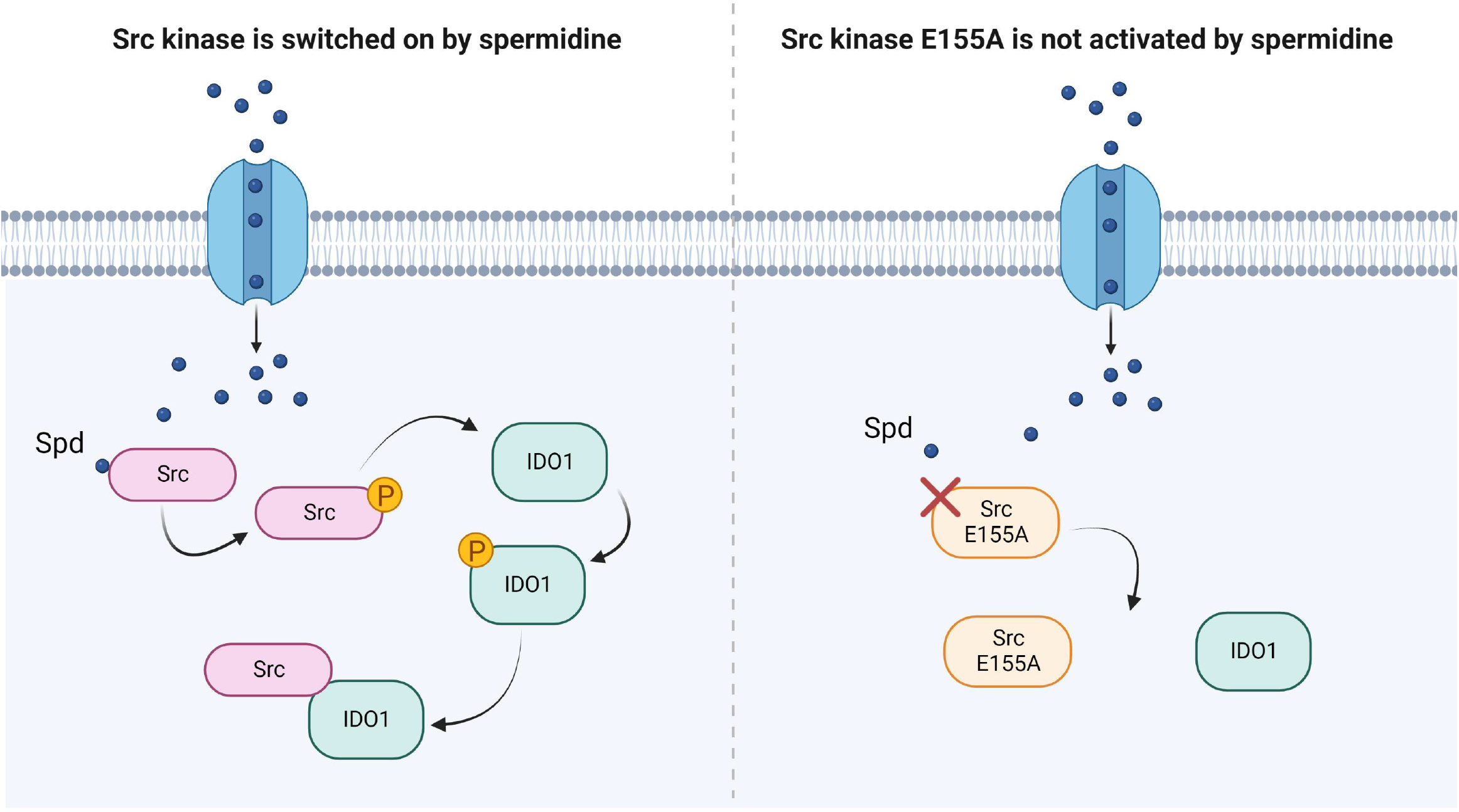
Scheme of the Src kinase modulation by the polyamine spermidine. Created with BioRender.com

## Materials and methods

### Cell lines and reagents

SYF cells (i.e., fibroblast null for Src, Fyn and Yes kinases (33); RRID:CVCL_6461) were grown in DMEM supplemented with 10% FCS, at 37 °C, in a humidified 5% CO2 incubator. Spermidine, LPA and recombinant Src protein were purchased from Sigma-Aldrich, while recombinant human IDO1 protein was obtained by Giotto Biotech. Construct expressing murine Src was obtained from Origene.

### Cell transfection and treatment

Src mutants were generated by PCR-based site-directed mutagenesis performed with overlapping and complementary primers containing the specific substitutions (**Table I**). The resulting PCR products were digested with appropriate restriction enzymes and cloned into pEF-BOS plasmid. Cells were transfected with 2 ug of the vectors expressing either wild-type Src or Src mutants according to the Lipofectamine 3000 protocol (Thermo Fisher Scientific). Stable transfectants were obtained by antibiotic selection of SYF cells transfected with pEF-BOS-based vectors

**Table I.**
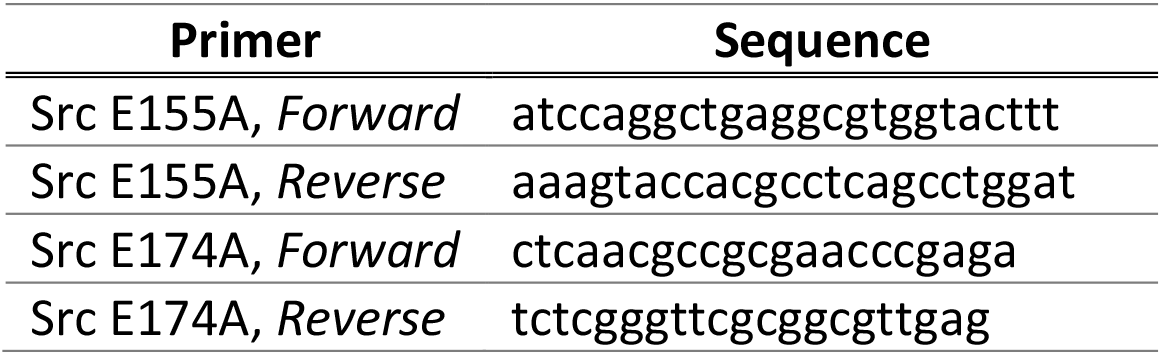
Primers for site-directed mutagenesis of Src.

carrying the puromycin resistant genes. SYF cells were serum-starved overnight before spermidine treatment. Cells were incubated with the polyamine or LPA for 60 minutes for immunoblot analysis, as otherwise indicated. These conditions were selected based upon optimization experiments.

### Split-luciferase fragment complementation assay

The N and C fragments of luciferase were amplified by PCR from pGL3-Basic. The fragment including nucleotide sequences of SH2 domain of Src (aa 374-465), linker (SRGGSTSGSGKPGSGEGSG), and Src consensus substrate peptide (WMEDYDYVHLQG), was synthesized by sequential reactions of PCR amplification. This cassette and the luciferase fragments were cloned into pCDNA 3.1 vector. SYF cells stably expressing the reporter were transfected with wild-type Src and Src E155A or Src E174A and then cultured into 96-w plate, in serum-free medium. After treatment with spermidine for 2h, cells were washed with PBS1X and lysed with PLB-lysis buffer. Luciferase activity was measured with the Luciferase Reporter Assay Kit (Promega).

### Immunoblot and co-immunoprecipitation studies

For immunoblotting, proteins were extracted in M-PER buffer (Thermo Fisher Scientific) supplemented with phosphatases and proteases inhibitors cocktails (Thermo Fisher Scientific) and run on SDS/PAGE. The pSrc/Src ratio was assessed with a rabbit Phospho-Src Family (Tyr416) Antibody (#2101, Cell Signaling Technology, Danvers, MA, USA; RRID:AB_331697), recognizing the phosphorylation at tyrosine 424 in murine Src, followed by the detection of total Src by rabbit monoclonal antibody (36D10, Cell Signaling Technology, Danvers, MA, United States; RRID:AB_2106059), as previously shown (27).

Co-immunoprecipitation appraises were performed following the manufacturer’s protocol (ThermoFisher) and as previously shown (51). Briefly, lysates were incubated over-night at 4°C with Dynabeads Protein G, prepared by blocking 12.5 µl of magnetic beads with PBS1X containing 0.5% BSA (w/v) and bound to 2.5 µg of rabbit anti-Src (36D10) or MultiMab™ Rabbit Phospho-Tyrosine (P-Tyr-1000; RRID:AB_2687925) antibody. After washing with buffer (25 mM citric acid, 50 mM Dibasic Sodium Phosphate dodecahydrate pH 5), the immuno-complex was eluted with Elution buffer (0.1 M Sodium Citrate dihydate pH 2-3) and Laemmli buffer. Proteins were run on SDS-PAGE and the expression of IDO1 was analyzed with a mouse anti-IDO1 antibody (clone 8G-11, Merck). Mouse monoclonal Ab against β-tubulin (Sigma-Aldrich; RRID:AB_2827403) was used as normalizer. Protein expression was measured by using Image Lab software (Bio-Rad) and the densitometric analysis of the specific signals was performed as previously described (31).

### Biochemical assay

For the in vitro cell-free assay, 5 ng of recombinant hSrc were combined with 10 μM of ATP and 100 μM of synthetic peptide (KVEKIGEGTYGVVYK) corresponding to amino acids 6-20 of p34cdc2. The reaction was carried out in a buffer containing 100 mM of Tris-HCl (pH 7.2), 125 mM of MgCl2, 25 mM of MnCl2, 250 μM of Na3VO2 and 2 mM of DTT. The mixture was incubated at 25°C for 30 minutes and the production of ADP was measured using the ADP-Glo kinase assay kit (Promega). Vmax was calculated after fitting the kinase activity data to the Michaelis–Menten equation. For the continuous in vitro assay, 50 ng of recombinant hSrc were incubated in the assay buffer with 300 ng of recombinant hIDO1, 100 μM of ATP, with or without spermidine (50 nM). The reaction was carried out at 25°C for the indicated time and then stopped by the addition of Laemmli buffer. Samples were run on SDS/PAGE and analyzed for the expression of Phospho-Tyrosine and IDO1 using an anti-pTyr -1000 and anti-IDO1 (clone 10.1, Merck) antibodies, respectively.

### Proximity ligation assay (PLA)

SYF cells expressing WT Src or Src E155A or Src E174A were serum-starved, stimulated with spermidine, fixed for 20 minutes with 4% PFA, permeabilized for 10 minutes with Triton-X 0.1% in PBS1X and then blocked. Duolink Proximity Ligation Assay (#DUO92008, Sigma-Aldrich) was performed according to the manufacturer’s protocol. Briefly, primary antibodies rabbit a-mouse Src (Thermo Fisher, 7G6M9) and mouse a-mouse IDO1 (clone 8G11, Merk) were conjugated with either PLUS (#DUO92009, Sigma-Aldrich) or MINUS (#DUO92010, Sigma-Aldrich) oligonucleotides to create PLA probes. Samples were incubated overnight at 4° C and, subsequently, ligase solution was added for 30 minutes. The signal was amplified with amplification polymerase solution at 37° C for 100 minutes. Nuclei were counterstained with 4’, 6’-diamidino-2-phenylindole (DAPI) (#DUO82040, Sigma-Aldrich). A total of 7 images (on average of 60 cells) per samples were taken with a Nikon inverted microscope (60X magnification) and analyzed with the software ImageJ.

### Electrostatic potential calculation study and docking study

The NMR structure of the SH2 domain of Src kinase (PDB ID: 2JYQ)(65) was taken from the protein data bank (www.rcsb.org) (66). Atomic coordinates were processed using the program PDB2PQR (67,68). The Adaptive Poisson-Boltzmann Solver (APBS) was applied to calculate the electrostatic potential of SH2 domain and map it on the excluded solvent surface (69). Specifically, the PARSE force field was employed with default parameters including a solute dielectric value = 2, solvent dielectric value = 78.54, solvent probe radius = 1.4 Å, and temperature = 298.150 °K. Two calculations were performed using cubic spline charge discretization and a grid dimension of 129 × 129 × 129 Å. The first run adopted a grid spacing of 0.574 × 0.557 × 0.509 Å, for a grid length of 73.433 × 71.279 × 65.163 Å centered at point 2.638 (x), 0.458 (y), 0.467 (z). The second run used a grid spacing of 0.494 × 0.484 × 0.456 Å for a grid length of 63.196 × 61.929 × 58.331 Å centered at the same point of the first run. The chemical structure of spermidine was taken from PubChem compound (70). The structure was processed using LigPrep (Schrödinger Release 2021-3: LigPrep, LLC, New York, NY, 2021) and applying the default settings.

The structure of the SH2 domain of Src kinase (PDB ID: 2JYQ) was processed employing the Protein Preparation Wizard (PPW) tool, as implemented in Maestro (Schrödinger Release 2021-3: Maestro, Schrödinger, LLC, New York, NY, 2021). In particular, hydrogen atoms were added and the internal geometries of the protein were optimized with a coordinate displacement restrain on heavy atoms set to 0.3 Å. The docking study was carried out defining a grid box for calculations centered on the center of mass of residues E155 and E174 (E4 and E23 according to 2JYQ sequence numbering). The inner box was sized 10×10×10 Å. Since the allosteric site features a shallow surface, a ligand induced-fit approach was used to investigate the binding mode of spermidine. Accordingly, docking solutions were obtained using the induced-fit docking algorithm (Schrödinger Release 2021-3: Induced Fit Docking, Schrödinger, LLC, New York, NY, 2021) and the standard protocol to generate up to n.20 binding poses of spermidine into the allosteric site. During calculations, ligand and receptor van der Waals scaling factors were set to 0.5 kcal/mol, respectively. The side chain conformations of residues within 5 Å of the ligand binding pose were sampled and refined using the OPLS 2005 force field. The structure of spermidine was then redocked with glide and standard precision (SP) scoring function into different obtained conformations of the allosteric site, using up to n.20 top energy conformations of the binding site within 30 kcal/mol of the minimum energy conformation.

## Acknowledgments

This research was funded by University of Perugia, Ricerca di base 2019, (RBGMON19; to G.M.), Associazione Italiana per la Ricerca sul Cancro (AIRC 2019-23084; to CV) and University of Perugia, Ricerca di base 2020 (INTEGRATE; to AM).

## Author contributions

Investigation, S.R., E.B., S.A, G.S., E.P., C.V.; Methodology, M.G., A.M., G.M.; Conceptualization, G.M. and M.G.; Supervision, F.F., A.M and C.O.; Writing-original draft preparation, G.M. All authors have read and agreed to the published version of the manuscript.

## Supporting information

Supplementary Figure S1. Spermidine does not modify neither the efficacy nor the affinity of Src kinase in the presence of increasing concentration of ATP.

Supplementary Figure S2. The glutamate residues E155 and E174 are conserved across different species.

Supplementary Table S1. Solutions of the docking study of spermidine into the allosteric site of Src SH2 domain.

Supplementary Figure S3. Efficient reconstitution of SYF cells with vectors coding for Src kinase.

## Data availability statement

All data generated or analyzed during this study are included in the manuscript and supporting file. Figure 1 - Source Data 1; Figure 2 - Source Data 2; Figure 3 - Source Data 3; Figure 3 - Source Data 4; Figure 3 - Source Data 5; Figure S3 - Source Data 6; Figure S3 - Source Data 7: contain the original blots used to generate the figures.

## Figure legends of Source data files

**Figure 1 - Source Data 1**. Original immunoblots of phosphorylated (pSrc), total Src and actin protein levels evaluated in cell lysates from SYF cells reconstituted with vector coding for wild-type Src and then treated with increasing concentration of spermidine.

**Figure 2 - Source Data 2**. Original immunoblots of phosphorylated (pSrc), total Src and actin protein level evaluated in cell lysates from SYF cells either reconstituted with vector coding for wild-type Src (WT) or Src mutated at glutamate 155 or 174 with alanine (mutants). Cells were then exposed to increasing concentration of spermidine.

**Figure 3 - Source Data 3**. Original immunoblots of immunoprecipitation with anti-phosphotyrosine antibody (IP) followed by the detection of IDO1 and Src with specific antibodies. Whole-cell lysates (PRE-IP) was used as control of protein expression of IDO1 and Src. SYF cells reconstituted with vectors coding for Src and IDO1 and then treated with spermidine (100 μM) for 60 minutes as well as cells transfected with vectors coding for either Src or IDO1 were used for the experiments.

**Figure 3 - Source Data 4**. Original immunoblots of in vitro kinase assay with rhIDO1 (300 ng) and rhSrc (50 ng) followed by immunoblot analysis with anti-phosphotyrosine and anti-IDO1 specific antibodies. The reaction was in either the presence or absence of spermidine.

**Figure 3 - Source Data 5**. Original immunoblots of immunoprecipitation of Src from SYF cells reconstituted with Src and IDO1, and then treated with spermidine for the indicated time. The detection of IDO1 and Src was performed by sequential immunoblotting with specific antibodies (IP). Whole-cell lysates (PRE-IP) was used as control of protein expression.

**Figure S3 - Source Data 6**. Original immunoblots of total Src and actin protein levels in cell lysates from SYF cells either reconstituted with vector coding for wild-type Src (WT) or Src mutated at glutamate 155 or 174 with alanine. SYF cells transfected with empty vector (SYF) were used as control.

**Figure S3 - Source Data 7**. Original immunoblots of phosphorylated (pSrc), total Src and β-tubulin protein level in cell lysates from SYF cells either reconstituted with vector coding for wild-type Src (WT) or Src mutated at glutamate 155 or 174 with alanine and then exposed to LPA (20 μM).

## Supplementary

**Supplementary Figure S1.**
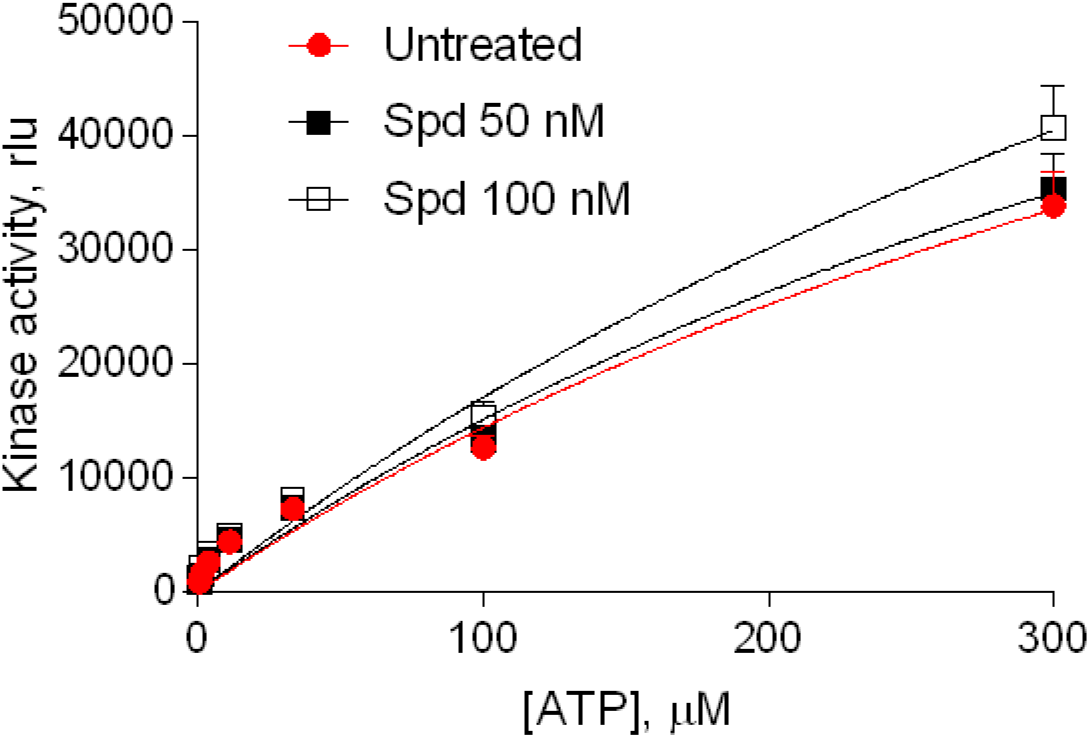
Spermidine does not modify neither the efficacy nor the affinity of Src kinase in the presence of increasing concentration of ATP. Enzymatic activity of recombinant human Src in the presence of fixed concentration of spermidine and increasing concentration of ATP. Vmax was calculated after fitting the kinase activity data to the Michaelis–Menten equation. Data were analyzed with one-way ANOVA followed by post-hoc Bonferroni test.

**Supplementary Figure S2.**
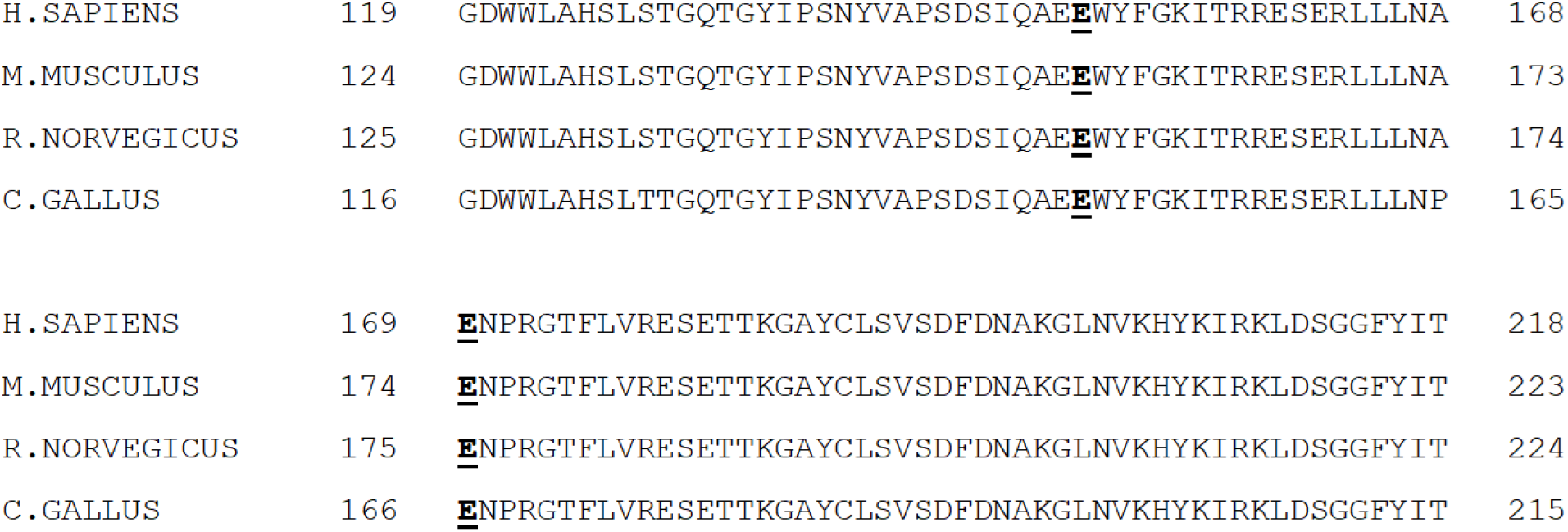
The glutamate residues E155 and E174 are conserved across different species. Alignment of the amino acid sequences of Src restricted to the stretch of amino acids containing the putative polyamine allosteric site. Conserved glutamate residues are highlighted in bold.

**Supplementary Table S1.** Solutions of the docking study of spermidine into the allosteric site of Src SH2 domain.

| Pose # | IFD Score (kcal/mol) | $\Delta$ IFD Score (kcal/mol) |
| --- | --- | --- |
| 1 | -227.50 | 0.00 |
| 2 | -227.41 | +0.08 |
| 3 | -227.11 | +0.39 |
| 4 | -227.03 | +0.47 |
| 5 | -227.00 | +0.50 |
| 6 | -226.95 | +0.55 |
| 7 | -226.70 | +0.79 |
| 8 | -226.68 | +0.81 |
| 9 | -226.68 | +0.82 |
| 10 | -226.66 | +0.83 |
| 11 | -226.58 | +0.92 |
| 12 | -226.56 | +0.94 |
| 13 | -226.54 | +0.95 |
| 14 | -226.49 | +1.01 |
| 15 | -226.43 | +1.07 |
| 16 | -226.34 | +1.15 |
| 17 | -226.15 | +1.34 |
| 18 | -225.31 | +2.19 |

**Supplementary Figure S3.**
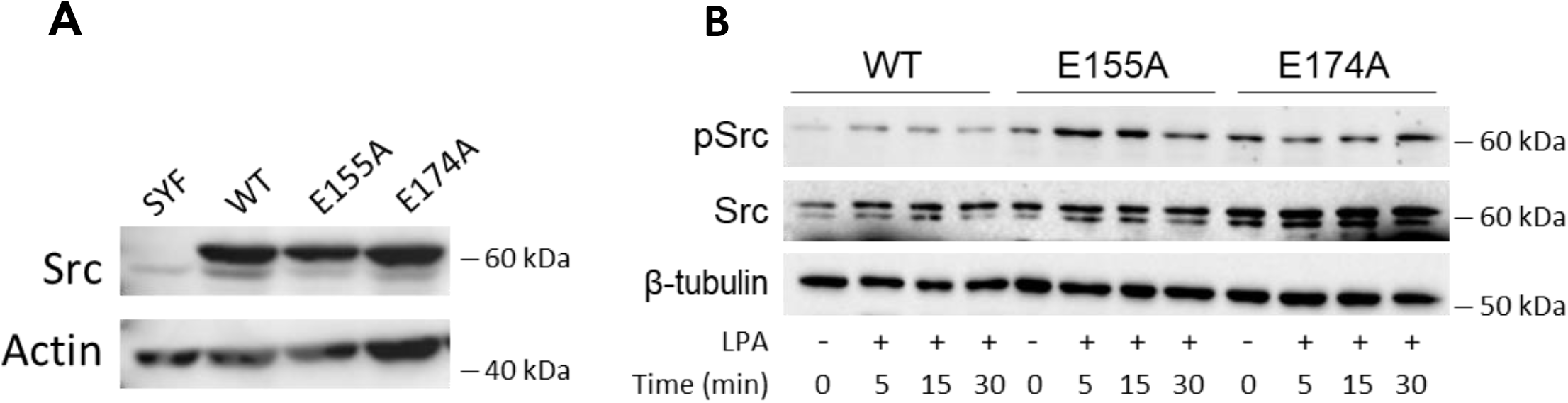
Efficient reconstitution of SYF cells with vectors coding for Src kinase. (**A**) Immunoblot analysis of total Src protein level in cell lysates from SYF cells either reconstituted with vector coding for wild-type Src (WT) or Src mutated at glutamate 155 or 174 with alanine (E155A; E174A). SYF cells transfected with empty vector (SYF) were used as control. Actin expression was used as normalizer. (**B**) Immunoblot analysis of phosphorylated (pSrc) and total Src protein level in cell lysates from SYF cells either reconstituted with vector coding for wild-type Src (WT) or Src mutated at glutamate 155 or 174 with alanine (E155A; E174A). Cells were then exposed to LPA (20 μM) for the indicated time. β-tubulin expression was used as normalizer.

